# NPmatch: Latent Batch Effects Correction of Omics data by Nearest-Pair Matching

**DOI:** 10.1101/2024.04.29.591524

**Authors:** Antonino Zito, Axel Martinelli, Mauro Masiero, Murat Akhmedov, Ivo Kwee

## Abstract

**Motivation:** Batch effects (BEs) are a predominant source of noise in omics data and often mask real biological signals. BEs remain common in existing datasets. Current methods for BE correction mostly rely on specific assumptions or complex models, and may not detect and adjust BEs adequately, impacting downstream analysis and discovery power. To address these challenges we developed NPmatch, a nearest-neighbor matching-based method that adjusts BEs satisfactorily and outperforms current methods in a wide range of datasets.

**Results:** We assessed distinct metrics and graphical readouts, and compared our method to commonly used BE correction methods. NPmatch demonstrates overall superior performance in correcting for BEs while preserving biological differences than existing methods. Altogether, our method proves to be a valuable BE correction approach to maximize discovery in biomedical research, with applicability in clinical research where latent BEs are often dominant.

**Data availability and implementation:** NPmatch is freely available on Github (https://github.com/bigomics/NPmatch) and on Omics Playground (https://bigomics.ch/omics-playground). The datasets underlying this article are the following: GSE120099, GSE82177, GSE162760, GSE171343, GSE153380, GSE163214, GSE182440, GSE163857, GSE117970, GSE173078, GSE10846. All these datasets are publicly available and can be freely accessed on the Gene Expression Omnibus (GEO) repository.

## 1. Introduction

Modern biomedical research employs high-throughput assays to generate single- and multi-omics data. For instance, RNA-sequencing data provides expression profiles of thousands of genes at genome-wide scale. Various experimental protocols at increasing granularity, including single-cell genomics, proteomics, or spatial transcriptomics have been developed and made accessible. Yet, bulk RNA-seq continues to be a widely used assay in current research practices.

However, these advancements are accompanied by significant challenges. One such challenge is the high cost of sample collection, processing and data generation, especially in studies involving a large number of samples (e.g., population-scale studies of disease). In large-scale studies, it is common practice to distribute the several steps of the data acquisition workflow across multiple centers. This often leads to the utilization of diverse protocols and technologies across the different centers. Additionally, research is increasingly relying on published datasets. Free, publicly available repositories like the Gene Expression Omnibus (GEO) database (Barrett et al., 2005), serve as valuable resources to scientists, offering quick access to existing datasets for re-analysis and to complement newly generated datasets.

Measurements in datasets generated in multiple centers will inevitably be affected by multiple sources of technical variation, collectively known as ‘Batch Effects’ (BEs). BEs may also arise within a single laboratory, due to distinct sequencing runs, depths, use of different sample donors, or when processing occurs in separate days. Cumulative variation can be also caused by smaller, hidden technical sources inherent to experimental settings. Altogether, BEs form a predominant, unwanted source of noise in omics data. BEs impact data mean and variance, and may confound real, underlying biological signal, altering false positive and false negative rates in downstream analyses e.g., (Kupfer et al., 2012, Tung et al., 2017, Johnson et al., 2007, Leek et al., 2010, Phua et al., 2022, Cuklina et al., 2021). Differential gene expression (DGE) testing, as an example, may be affected by BEs. This is especially true in cases where the variable of interest is highly unbalanced between distinct batches. To minimize BEs, it’s crucial for the study design to involve a balanced representation of samples across batches. Unfortunately, study designs are often imperfect. When the variable of interest is highly imbalanced between distinct batches, it can become very challenging to disentangle biological signals from BEs.

Previous studies have assessed the extent to which BEs impact measurements and discovery power e.g., (Leek et al., 2010, Leigh et al., 2018, Lauss et al., 2013, Rasnic et al., 2019). Particularly in large datasets, BEs may underlie inconsistencies across studies. To address BEs computationally, batch correction methods have been developed. On a general level, these can be categorized into (i) ‘Supervised methods’ such as ComBat (Johnson et al., 2007) and Limma’s RemoveBatchEffects (Ritchie et al., 2015), which use linear models to adjust known batch effects; (ii) ‘Unsupervised methods’, such as SVA (Leek and Storey, 2007) and RUV (Gagnon-Bartsch and Speed, 2012), which attempt to identify potential sources of variation due to BEs without requiring prior knowledge of the batch vector. These methods mostly rely on complex assumptions or models, and would thus need approximated distributions with uncertain distortion from the model-expected distribution. Furthermore, batch correction methods suffer from the inherent heterogeneity both within and between batches, which is exacerbated in an unbalanced mix between study groups and batches in the absence of matching replicates between batches. As a result, they may not necessarily detect or adjust BEs adequately and consistently across diverse datasets. In order to achieve an unbiased BE correction, both batch and phenotype labels would be needed. While this may be the case for fully controlled experiments, it’s unrealistic in clinical research where BEs are often unknown and phenotype classes of patient biopsies are often undefined.

Here, we developed a batch correction method, NPmatch (nearest-pair matching), that relies on distance-based matching to deterministically search for nearest neighbors with opposite labels, so-called “nearest-pair”, among samples [Fig. 1A-C; Methods]. NPmatch requires knowledge of the phenotypes but not of the batch assignment. Differently to many other algorithms, NPmatch does not rely on specific models or underlying distribution. It does not require special experimental designs, randomized controlled experiments, control genes or batch information. NPmatch is based on the simple rationale that samples should empirically pair based on distance in biological profiles, such as transcriptomics profiles.

**Figure 1.**
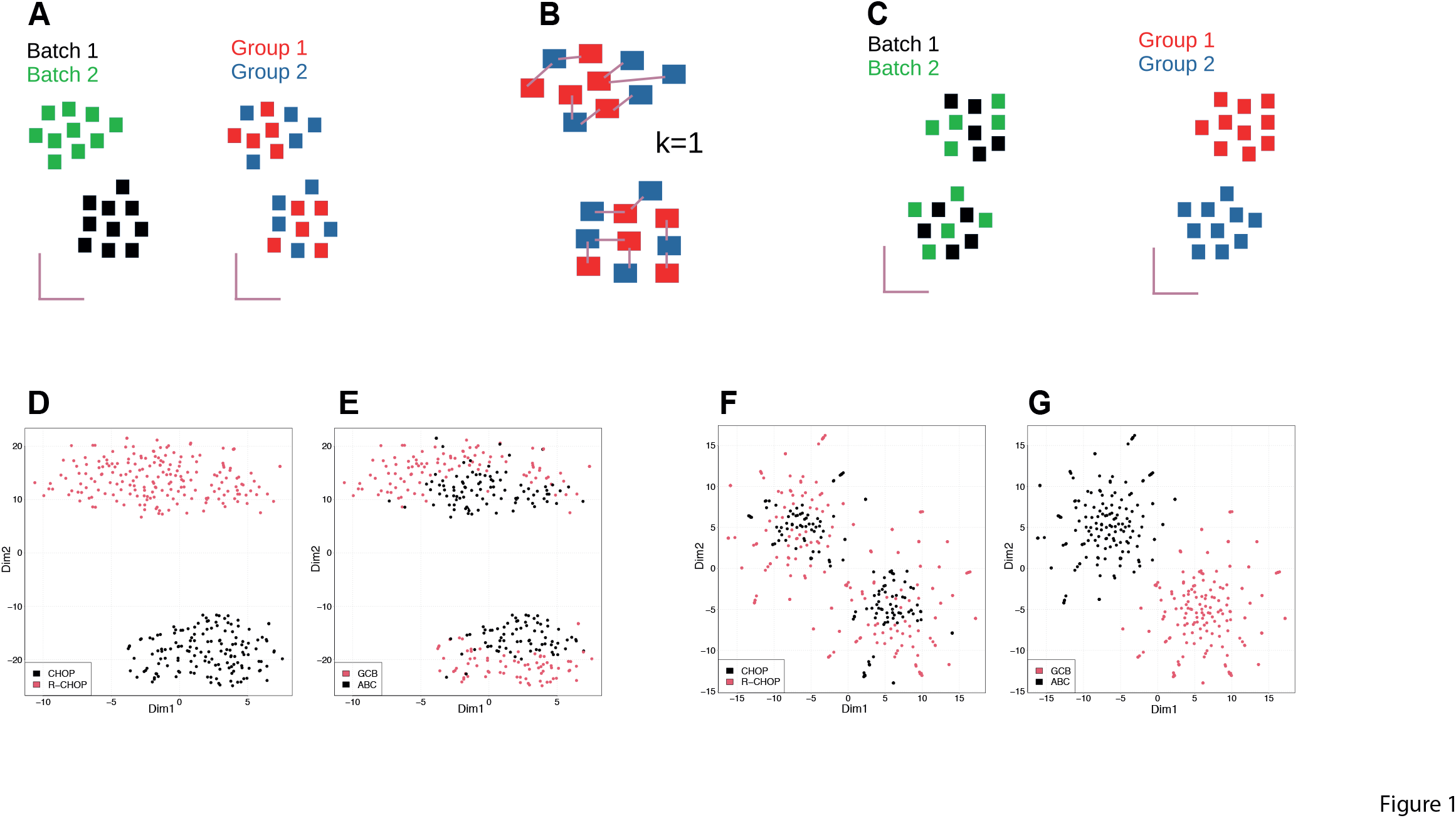
NPmatch algorithm and testing on a batched dataset. (A) Representative clustering of a dataset affected by batch effects. (A) Samples segregate by batches rather than biological group. (B) NPmatch conducts nearest neighbor search for each sample (Methods). A k=1 has been chosen as representative illustration. (C) NPmatch results into a batch-corrected dataset, where samples segregate by biological condition of interest rather than batches. t-Distributed Stochastic Neighbor Embeddings (t-SNE) projections on the first two dimensions of batched data (D-E) and (F-G) batch-corrected data in a real dataset (GSE10846; Methods).

Our method was inspired by principles of the statistical matching theory (M. D’Orazio, 2006). Distinct matching methods have been made available through integrated frameworks. One is ‘*MatchIt*’ (Ho, 2007, Daniel Ho, 2011), which performs matching as a form of subset selection with pruning and weighting. Similarly, our method performs unit (sample) selection to classify the units into the distinct phenotype groups, and then performs nearest neighbor search (NNS) through correlation or Euclidean distance between units. As NPmatch uses prior knowledge on phenotypic groups, it relies on a form of data stratification. Similarly, matching may also involves stratification, though with different modalities (Zubizarreta, 2014, Austin, 2014). The NNS results into pairs of units within and across condition classes. As NNS results into a fully weighted dataset (i.e., weight (distance) associated to each unit), the *k* closest units can be determined for each unit within each group. The NNS is nonparametric as it is neither based on propensity scores nor depends on regression parameters. Instead, it is based on sample distances within the stratified dataset, with pairs fully drawn from the original dataset. Different to original matching techniques, NPmatch enables full dataset matching: all available units are matched to *k* units in the group with opposite label. No units are dropped or removed. Our method generates a corrected data matrix where the unwanted effect (i.e., batch-related variables) is removed through linear regression in Limma. This results in a batch corrected dataset suitable for downstream analyses.

We conducted extensive testing of NPmatch in 11 publicly available microarray and RNA-seq datasets. We assessed multiple BE correction metrics, including number of differentially expressed genes between the conditions of interest, principal component analysis, silhouette score for clustering, and non-linear dimensional reductions. We demonstrate that NPmatch tackles BEs satisfactorily while preserving the biological heterogeneity between samples. Remarkably, NPmatch outperforms the commonly used batch correction methods Limma (Ritchie et al., 2015), ComBat (Johnson et al., 2007), SVA (Leek and Storey, 2007), RUV (Gagnon-Bartsch and Speed, 2012) and PCA (Giuliani, 2017, Jolliffe and Cadima, 2016).

## 2. Materials and Methods

### 2.1 NPmatch algorithm

The input to NPmatch is a normalized and log-transformed gene expression matrix X^p x n^ (p=features, n=samples), which may suffer from noticeable or latent batch effects. NPmatch does not require knowledge on the batches. Instead, NPmatch requires the phenotype vector. For a more efficient computation (beneficial for large datasets or when testing numerous datasets), NPmatch can select the top variable features (genes). The features are feature-centered and then further centered per condition group. Given X^p x n^, where n samples are distributed across *c* condition/phenotypic groups of biological interests, the rationale here is to buffer potentially significant differences in average expression between the two groups driven by or affecting the top genes. Inter-sample similarities are then determined by either computing the Pearson correlation matrix D^n x n^ (default) or Euclidean distance. For convenience, D is transformed into a 1-(D) scale such that both positive and anti-correlations are handled within the 0-1 range of values (0 highest correlation; 1 lowest correlation). The Pearson correlation matrix D^n x n^ is subsequently decomposed into the *c* phenotypic/condition groups. For each sample, a *k* nearest-neighbor like search is conducted to identify the closest k-nearest samples across each *c* phenotypic/condition group. The process results into a matrix X^n x (k x c)^ where for each sample, k-nearest samples are identified per each *c* condition. The X^n x (k x c)^ matrix is then used to derive a (i) vector of length L=n x k x c, storing all the computed pairs; (ii) a fully paired dataset X^p x L^. As pairing may *per-se* imply duplication of correlated signals (which is a BE-like effect), Limma ‘RemoveBatchEffect’ is used to correct for the ‘pairing effects’ through linear regression (Ritchie et al., 2005). The batch-corrected X1^p x L^ matrix is finally condensed into its original p x n size by computing, per each feature, the average values across duplicated samples. Thus, the X1^p x n^ matrix represents the batch-corrected dataset which can be used for further downstream analyses.

### 2.2 Datasets

NPmatch’s performance was tested on 11 publicly available human RNA-seq datasets (Sprang et al., 2022), and a microarray dataset, and compared to Limma (Ritchie et al., 2015), ComBat (Johnson et al., 2007), SVA (Leek and Storey, 2007), RUV (Gagnon-Bartsch and Speed, 2012) and PCA. All datasets had available expression data and batch information. A brief description of each dataset is provided below.

- GSE120099 (Lo Sardo et al., 2018): Induced Pluripotent stem cells were generated from individuals carrying the 9p21.3 risk locus for coronary artery disease, and from non-risk individuals. Genome editing was used to delete the haplotype, vascular smooth muscle cells (VSMCs) were generated and RNA-seq performed. Dataset for testing included a total of 92 samples (48 KO, 44 WT) split across 3 batches.
- GSE82177 (Wijetunga et al., 2017): RNA-seq from liver biopsies of 27 samples (10 uninfected controls, 9 HCV-infected non-tumor samples, 8 HCV-infected HCC tumor samples) split across 2 batches. Control samples and non-tumor samples were combined into a single group prior to batch effect assessment.
- GSE162760 (Farias Amorim et al., 2021): RNA-seq from whole blood samples from Leishmania braziliensis-infected individuals and non-infected controls. Dataset for testing included a total of 64 samples (14 non-infected controls, 50 Leishmania infected samples) split across 6 batches.
- GSE171343 (Bowles et al., 2021): Induced pluripotent stem cell-derived cerebral organoids expressing tau V337M mutation and CRISPR-corrected isogenic controls were generated and RNA-seq performed at distinct differentiation stages. Dataset for testing included a total of 240 samples (100 V337M, 140 V337V) split across 3 batches.
- GSE153380 (Alvarez-Benayas et al., 2021): RNA-seq was performed on 5 primary Plasma Cells (PC), 28 Multiple Mieloma (MM) PC, and 5 cell line samples. Samples ‘A26.19’ (PC) and ‘A27.22’ (PC) appeared to be merged with A26.18 (PC) and A27.21 (PC), respectively, at source. For testing we included a total of 26 samples (23 MM, 3 PC) split across 3 batches.
- GSE163214 (Procida et al., 2021): RNA-seq was performed on HeLa Kyoto cells following knockdown of *JAZF1* and control cell lines. The following two samples were removed as corresponding data appeared corrupted at source: ‘GSM4975193_siJAFZ1_Rep2_Batch1’ and ‘GSM4975199_siJAFZ1_Rep5_Batch2’. Dataset for testing included a total of 8 samples (5 controls, 3 KD) split across 2 batches.
- GSE182440 (Lim et al., 2021): RNA-seq was performed on postmortem putamen samples of control subjects and subjects affected with alcohol use disorder (AUD). Dataset for testing included a total of 24 samples (12 control, 12 AUD) split across 2 batches.
- GSE163857 (Moser et al., 2021): RNA-seq was performed from (i) microglia cells sorted from human-APOE carrying mice; (ii) microglia cells differentiated from human induced pluripotent stem cells from healthy subjects genotyped for APOE, untreated and treated with the heavy metals Cadmium (Cd) or Zinc (Zn). For testing we included the 24 human microglia samples (15 control, 4 Cd-treated, 5 Zn-treated) split across 2 batches.
- GSE117970 (Cassetta et al., 2019): RNA-seq of purified monocytes and tumor-associated macrophages from breast cancer biopsies, endometrial cancer biopsies, and normal tissues. For testing we included a total of 88 samples (50 normal, 38 breast cancer samples) split across 5 batches.
- GSE173078 (Kim et al., 2021): RNA-seq was performed from gingival tissue biopsies in states of periodontal health, gingivitis, and periodontitis disease. Dataset for testing included a total of 36 samples (12 healthy control, 12 gingivitis, 12 periodontitis) split across 2 batches.
- GSE10846 (Lenz et al., 2008): Array expression profiling was performed on clinical samples from diffuse large B-cell lymphoma (DLBCL) patients pre-treated with the chemotherapy regimens CHOP and Rituximab-CHOP. Dataset for testing included a total of 350 samples (167 ABC, 183 GCB) split across 2 batches (CHOP, R-CHOP).

### 2.3 Datasets preprocessing

All datasets were processed consistently within the same pipeline. For each dataset, the raw data were downloaded from GEO along with associated metadata and processed in

R. If feature (gene) identifiers were not official gene symbols, the official gene symbol was retrieved and assigned. In rare cases of duplicated gene symbols, the average expression values across duplicated features was calculated per sample and duplicated features removed. Genes undetected across all samples were removed. Expression data were normalized (i) within samples using counts per millions (CPM) followed by log2+1 transformation, and (ii) across-samples using quantile normalization in limma (Ritchie et al., 2005). Normalized data were used as input to the distinct batch correction algorithms.

### 2.4 Methods and Metrics for BEs detection and correction

The following methods and metrics were employed to assess BEs in the uncorrected datasets and upon batch correction:

- Silhouette score (SS): SS measures how well samples of the same group cluster together. SS values are defined within the range [-1,+1], where lower values indicate poor matching and clustering, and higher values indicating good match. Thus, BEs could be assessed with the SS, whereby higher values are expected upon batch correction. SS are computed using the R package ‘cluster’.
- Signal-to-Noise Ratio (SNR) of Log2FC: SNR is a well standardized measure in high-dimensional data, particularly genomic data. SNR measures the ratio between a signal of interest and a background noise in the underlying data. As signal, we utilize the average Fold-Change (FC) (in the Log2 scale) calculated through differential gene expression analyses (see below) between the phenotypes/condition of interests. The noise is defined as the average features’ standard deviation across all samples in the data matrix.
- PC1 Ratio: Singular value decomposition (svd) is applied to the data matrix. For each phenotype class, the absolute Pearson’s correlation between each singular value and the phenotype label is computed (across all samples). In order to assess the overall extent to which variation in the data may be due to phenotype, the average correlation across the phenotypes is then computed for each PC. We define PC1 Ratio as the ratio between the value of the first PC and the sum of the values of all available PCs. The higher the PC1 Ratio the better the batch correction.
- Differential Gene Expression (DGE) testing: In principle, one may expect that appropriate batch correction should improve the signal to detect biologically meaningful differences between phenotypes/condition of interests. This holds true both compared to uncorrected data (i.e., batch-confounded data) and data with inefficient batch adjustment. On the basis of this principle, DGE was performed between phenotypes/conditions of interests in both uncorrected data and upon batch correction using linear models and moderated t-test in limma. Differentially expressed genes (DGEs) are defined if absolute Log2FC≥0.5 and FDR≤0.05. Number of DEGs was used as a comparative metric between BE correction methods.

We sought to compute a score for each batch correction method. To this end we first computed the ratio between number of DEGs, SNR, and SS of the corrected data versus the uncorrected data matrix. As the uncorrected dataset was used as reference, the score is always 1 for the uncorrected data. The geometric mean of the ratios was then calculated as an integrated score of overall performance of each method in each dataset. To have a metric representative of overall method’s performance across all tested datasets, we computed the mean rank of the score for each method across all tested datasets.

## 3. Results and Discussion

BEs represent a major source of unwanted variation in high-dimensional data. BEs mask meaningful biological signals across conditions of interest and can impact discovery and reproducibility. In this work, we present NPmatch, a new method for BE correction [Fig. 1A-C; Methods]. It relies on the rationale that samples should empirically pair based on their distance in biological profiles. NPmatch is not restricted to prior assumptions on the nature of BEs. It also works in studies where the requirement of balanced sample distribution among batches is violated, which reasonably occurs due to the logistic and technical limitations in clinical research. We tested NPmatch in 11 microarray and RNA-seq datasets spanning diverse scenarios in terms of sample size and balanced representation of samples between batches, and compared to supervised and unsupervised methods, including limma ‘RemoveBatchEffects’, ComBat, SVA, RUV and PCA correction.

We initially tested NPmatch on a large batched array expression dataset of activated B-cell (ABC) and germinal center B-cell (GCB) diffuse large B-cell lymphoma (DLBCL) samples pre-treated with two different pharmacological regimens (Lenz et al., 2008). As treatment was performed prior to expression profiling and samples were split in the two groups for processing, this dataset well represent a scenario of how BEs may impact the data. In the uncorrected data BEs appear evident with samples clustering by pharmacological treatment [Fig. 1D-E]. NPmatch successfully corrects the BEs, with samples clustering by DLBCL type, reflecting their biological heterogeneity [Fig. 1F-G]. In another complex representative dataset (GSE162760; Methods), NPmatch achieves better batch correction while reasonably preserving the biological heterogeneity between samples compared to other methods [Fig. 2A]. The batch-corrected data demonstrate that samples part of the same phenotypic class cluster together. Assessment of t-SNE plots reveals that when compared to other methods, NPmatch demonstrates better clustering of samples based on the biological variable of interest, in most of the tested datasets [Fig. S1]. Accordingly, BEs appear substantially attenuated upon batch correction [Fig.S2]. As a control, we also performed batch correction with all methods upon randomization of the phenotype classes. As expected, no appropriate batch correction was achieved [Fig. S3].

**Figure 2.**
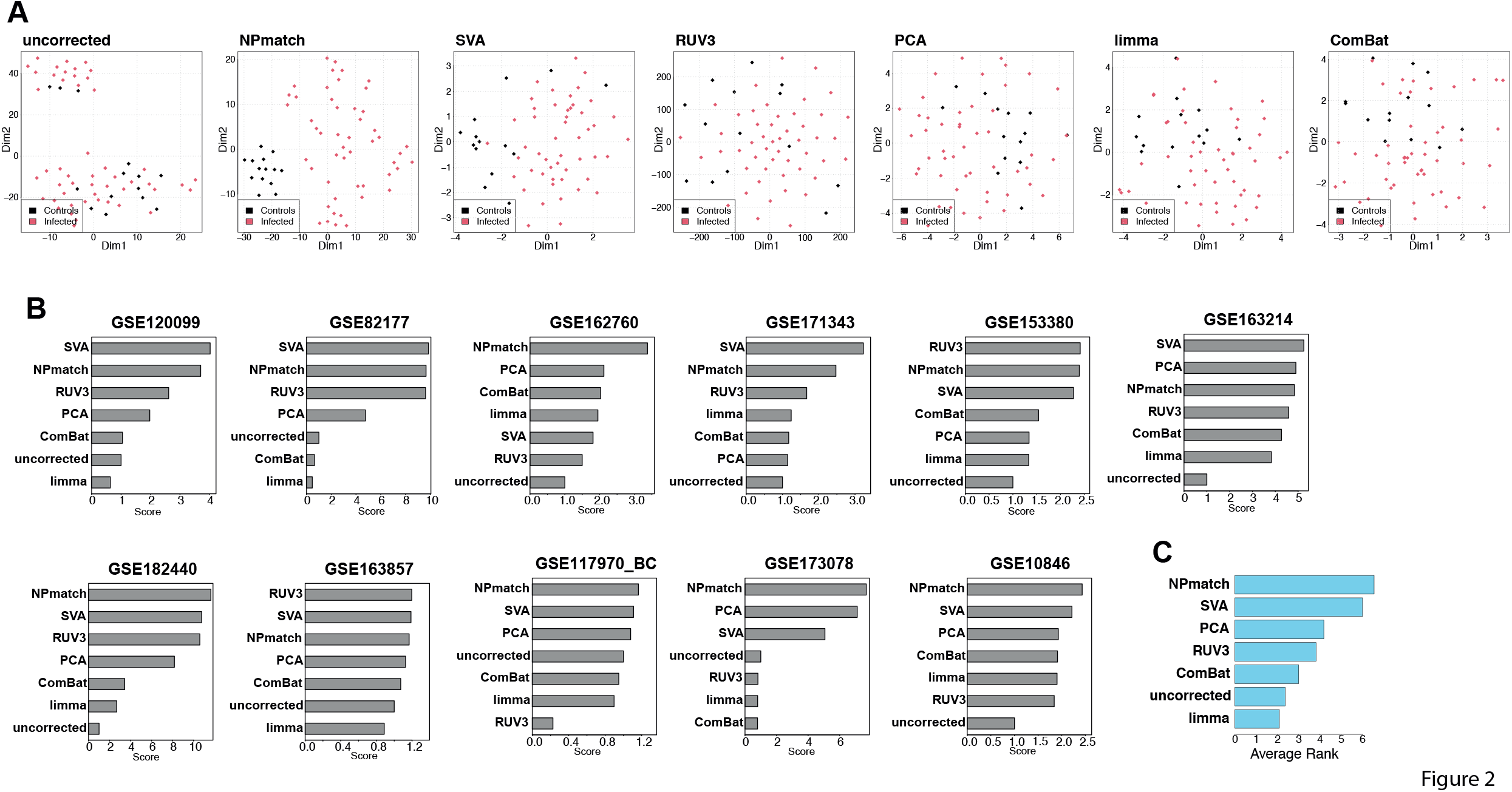
Comparison between NPmatch and other batch-correction methods. (A) t-SNE of uncorrected and batch-corrected data for GSE162760 (Methods). In each plot the samples are colored by the phenotype variable. The batch correction method employed is indicated at the top of each plot. (B) Bar plots of performance score (integrating multiple batch correction metrics; Methods) for each batch correction method in each tested dataset. (C) Ranked bar plot of mean rank score (Methods) for each BE correction method across all datasets.

To assess the extent to which batch correction impacts biologically meaningful signals in the data, we computed the number of differentially expressed genes between the conditions of interest, signal-to-noise ratio, silhouette score, and correlation between principal components and phenotype labels, in the uncorrected data and following batch correction. We found that NPmatch outperforms existing methods for most of the assessed metrics in the tested datasets [Fig.S4]. Likewise, upon combination of the metrics (Methods), NPmatch emerged among the top performing methods for the majority of datasets [Fig.2B]. We also computed an overall metric representative of each method’s performance across all datasets (Methods). In line with results from each single metric, NPmatch exhibited an overall superior performance than the other methods in most cases [Fig.2C].

Altogether, the data indicate that NPmatch tackles BEs satisfactorily while also preserving the biological heterogeneity between samples. This is proved by (i) the high number of DEGs detected and (ii) the improved clustering of samples in the dimensionally-reduced space. NPmatch also preserves the original distribution of the data. Remarkably, we also demonstrate that NPmatch outperforms or ranks among the top when compared to the highly used batch correction methods Limma, ComBat, SVA, RUV and PCA.

We applied NPmatch only to bulk transcriptomics data as the algorithm does not support single-cell level data. In fact, while NPmatch needs the phenotype labels (but not the batch labels), in single-cell RNA-seq data the phenotype labels - typically the cell types – are unknown and the batch information is usually available. Importantly, we believe NPmatch may also reasonably accommodate other high-dimensional, noisy data types, such as peptide and proteomic data. However, these data types are associated with other problems. For example, prior to batch correction for proteomics data, one should address the question of whether the preprocessing steps of normalization and imputation should be performed ahead of batch-correction (e.g., to avoid missing values) or upon appropriate data transformation and batch correction. Thus, applying NPmatch to other data types warrant separate studies.

While we recognize that there may not be a single, all-encompassing solution to address BEs in RNA-seq data or other biological data types given the inherent heterogeneity present in batched datasets, we propose NPmatch as a powerful alternative method, especially when other methods fail to resolve BEs.

## Acknowledgements

We thank the team at BigOmics Analytics for providing valuable feedback on this work.

## FIGURE LEGENDS

**Figure S1. t-SNE plots of uncorrected and batch corrected data to assess clustering based on phenotype labels.** The samples are colored by the biological variable of interest as per each dataset’s metadata. Dataset GEO identifier, batch correction method employed and score (Methods) are reported at the top of each plot.

**Figure S2. t-SNE plots of uncorrected and batch corrected data to assess clustering based on batch labels.** The samples are colored by the batch labels as per each dataset’s metadata. Dataset GEO identifier and batch correction method employed are reported at the top of each plot.

**Figure S3. t-SNE plots of uncorrected and batch corrected data to assess clustering following randomization of the phenotype labels.** The samples are colored by the phenotype labels as per each dataset’s metadata. Dataset GEO identifier and batch correction method employed are reported at the top of each plot.

**Figure S4. Assessment of BE correction metrics for each method in each tested dataset.** Bar plots show the number of differentially expressed genes (on the log2 scale) between the conditions of interest, signal-to-noise ratio, silhouette score, and correlation between principal components and phenotype labels (Methods), in the uncorrected data and following batch correction.

