## Supplementary figures and images for "NPmatch: Latent Batch Effects Correction of Omics data by Nearest-Pair Matching"

### Figure S1

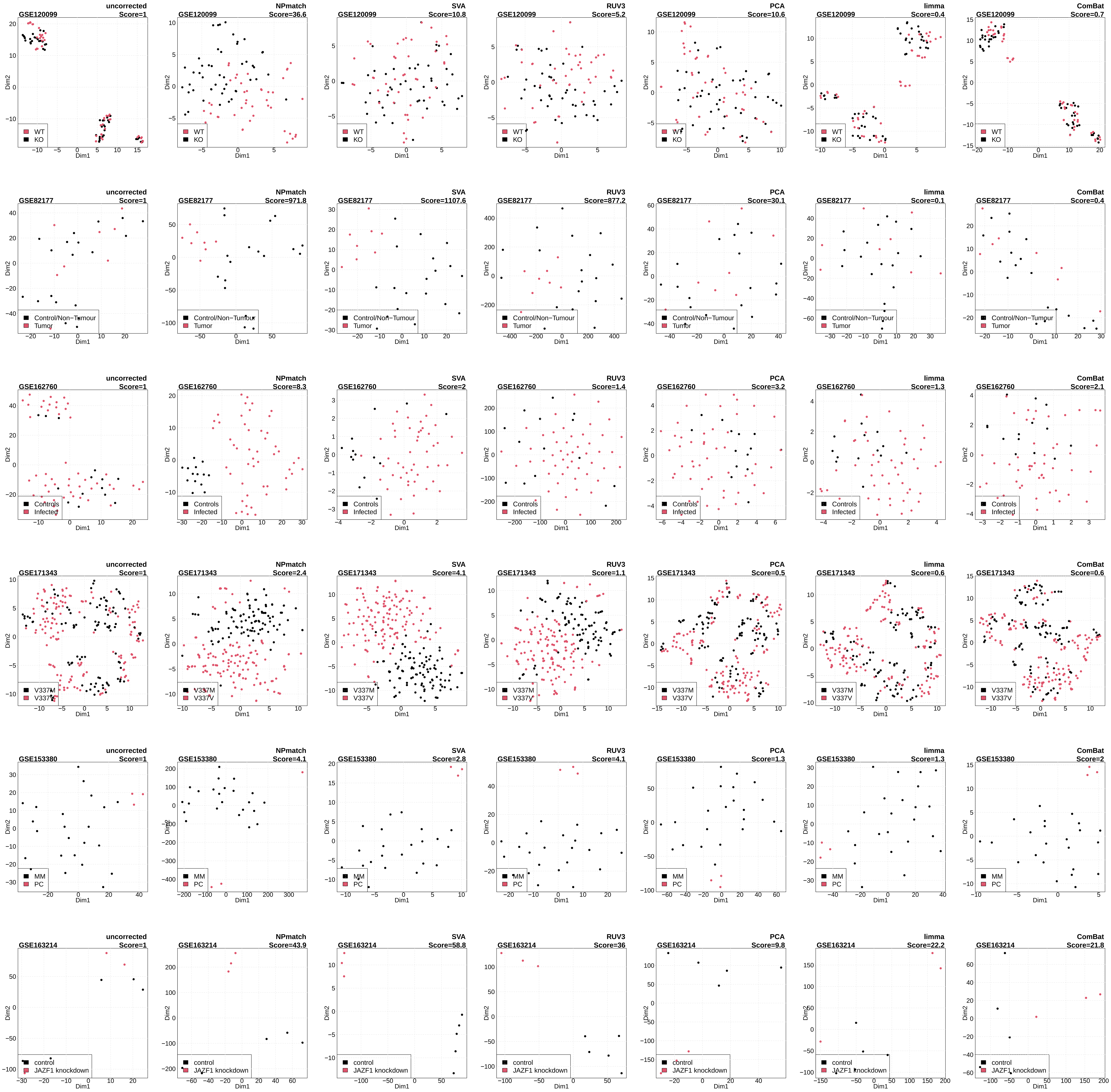

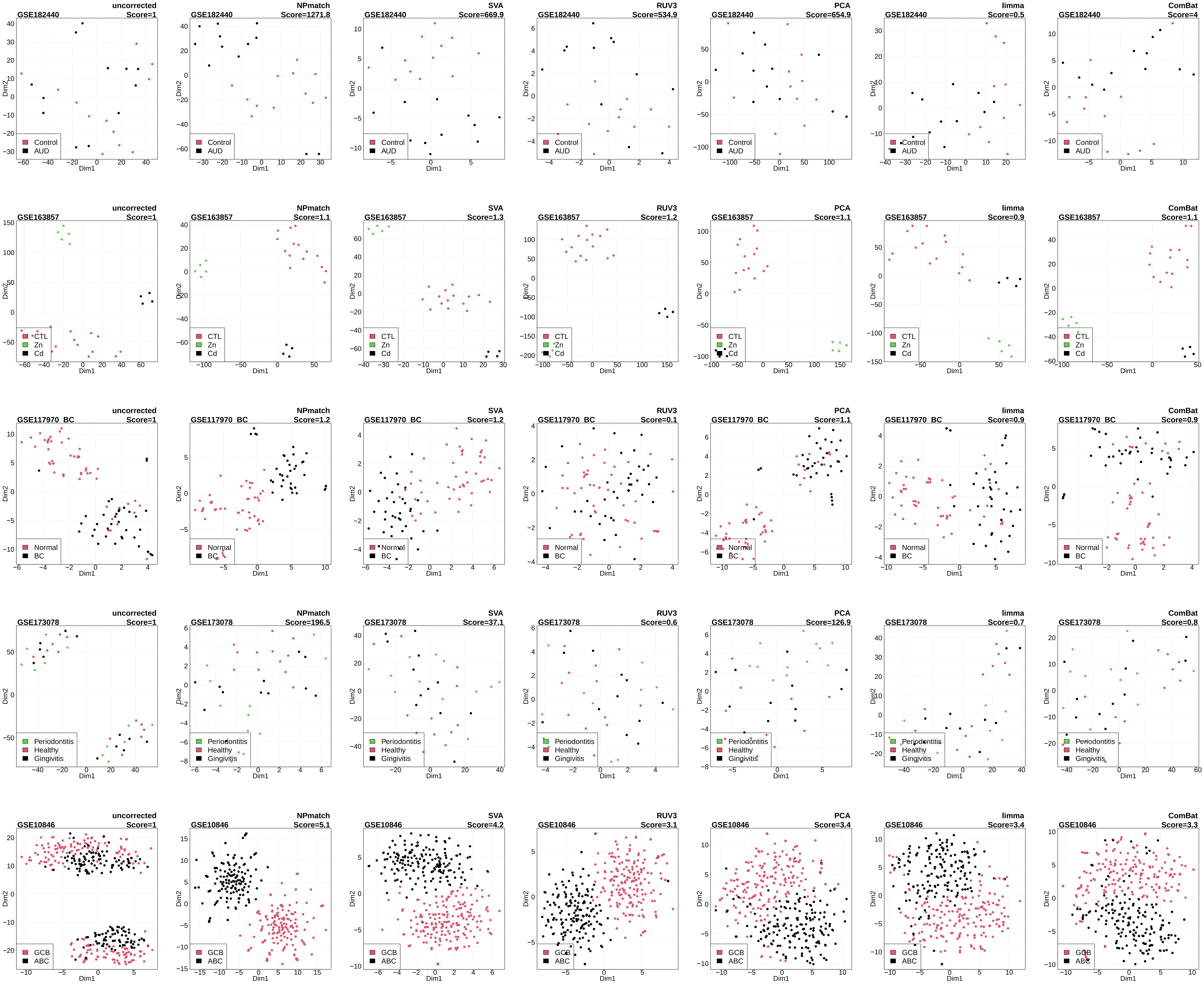

### Figure S2

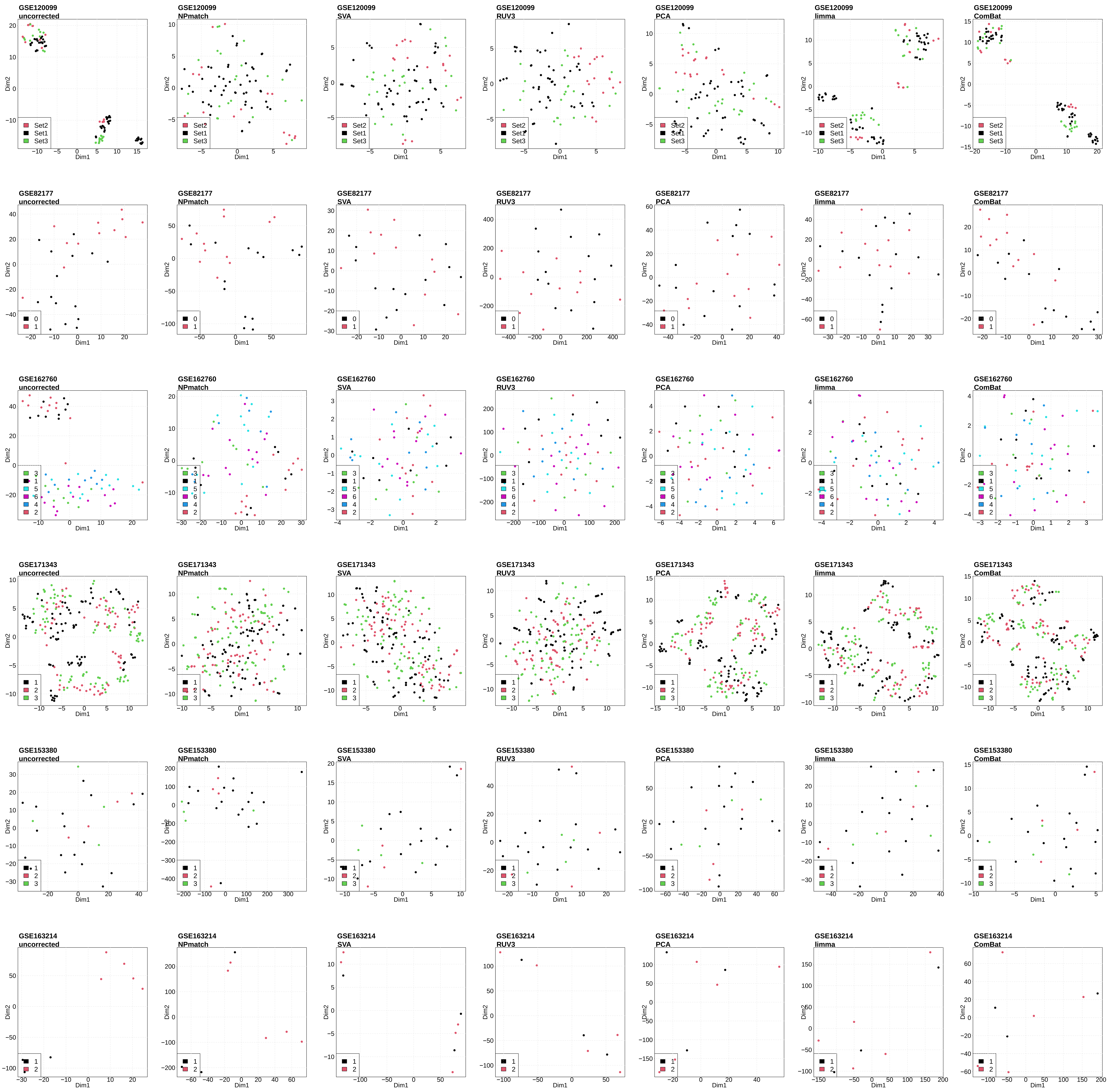

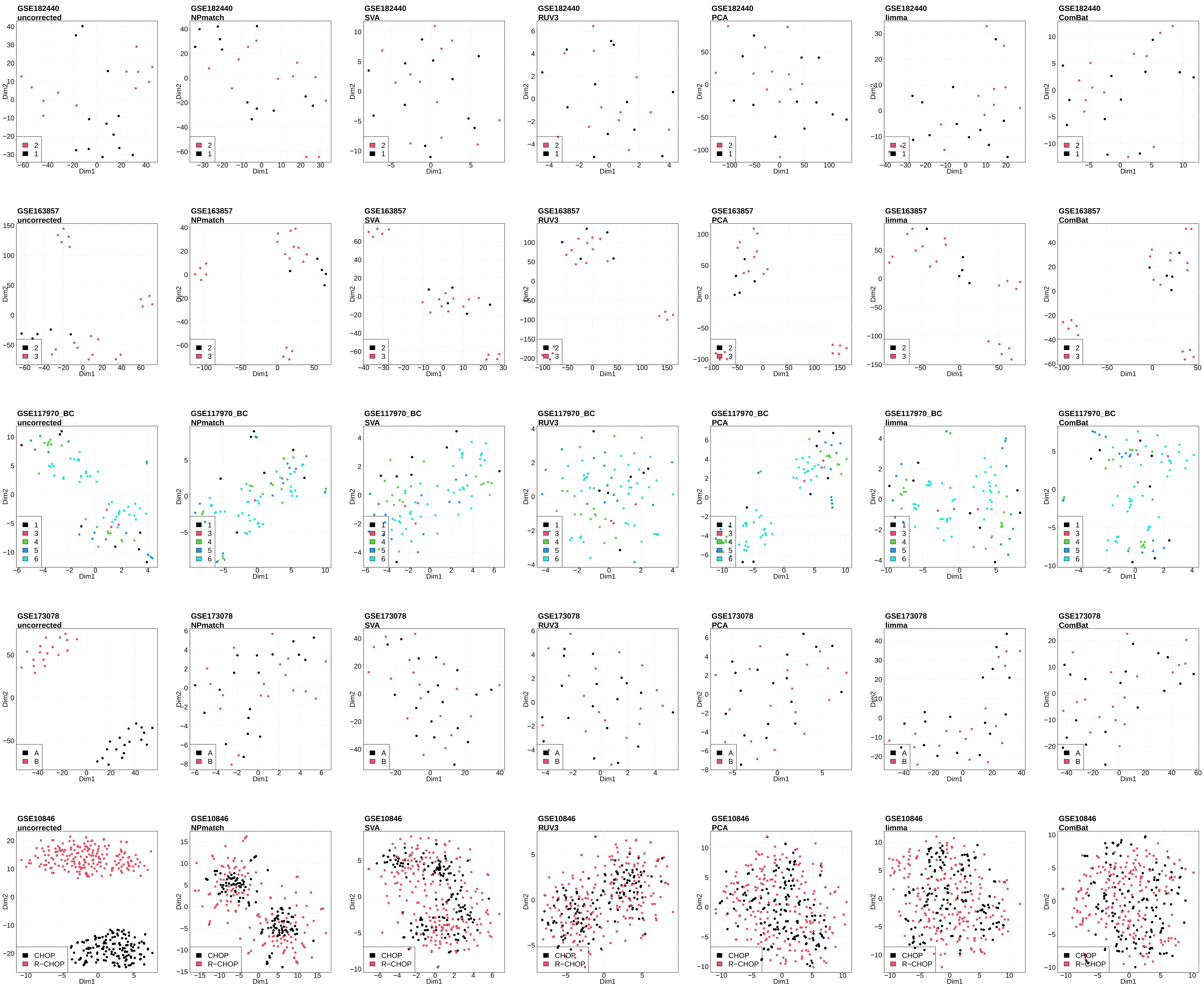

### Figure S3

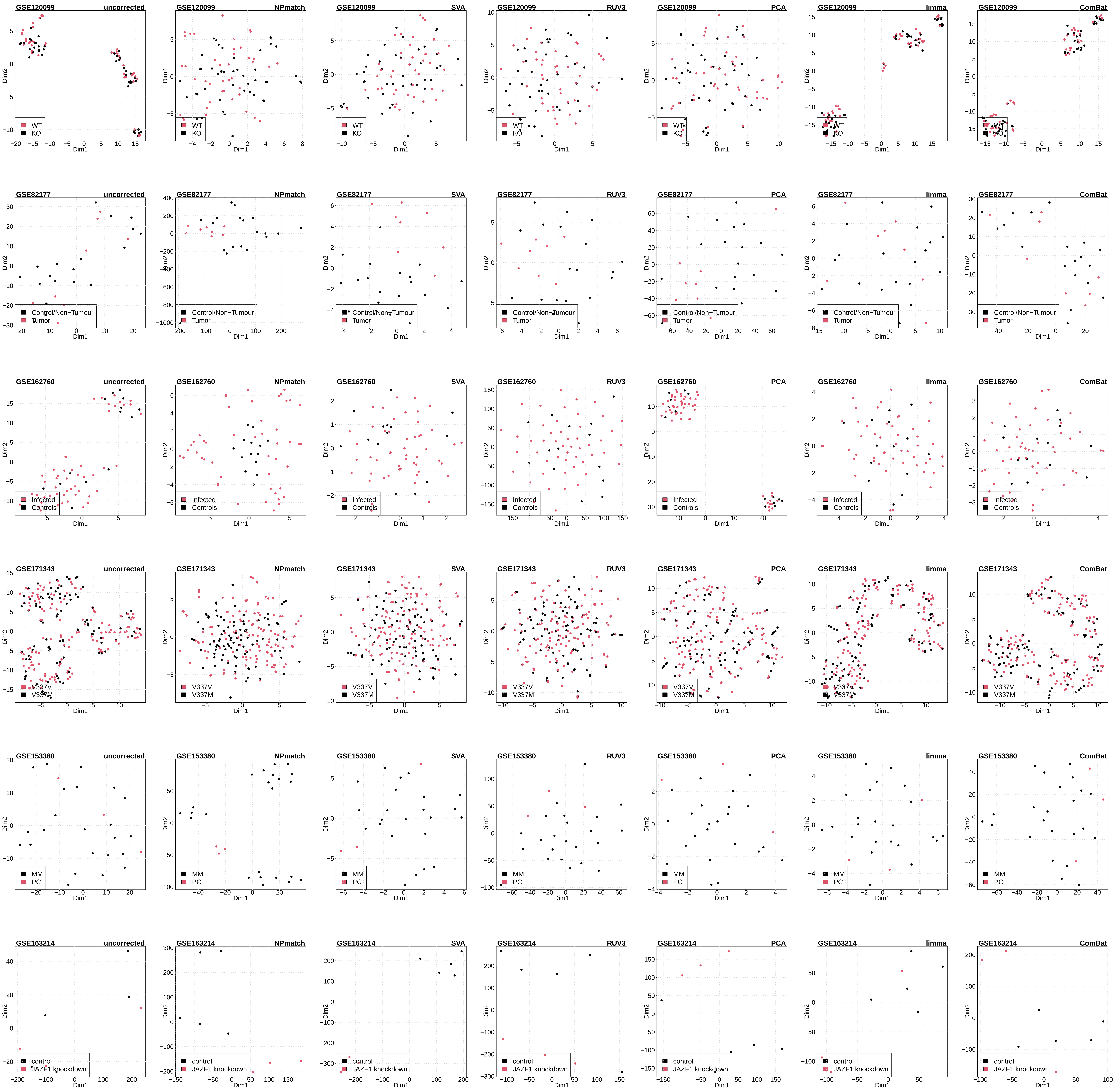

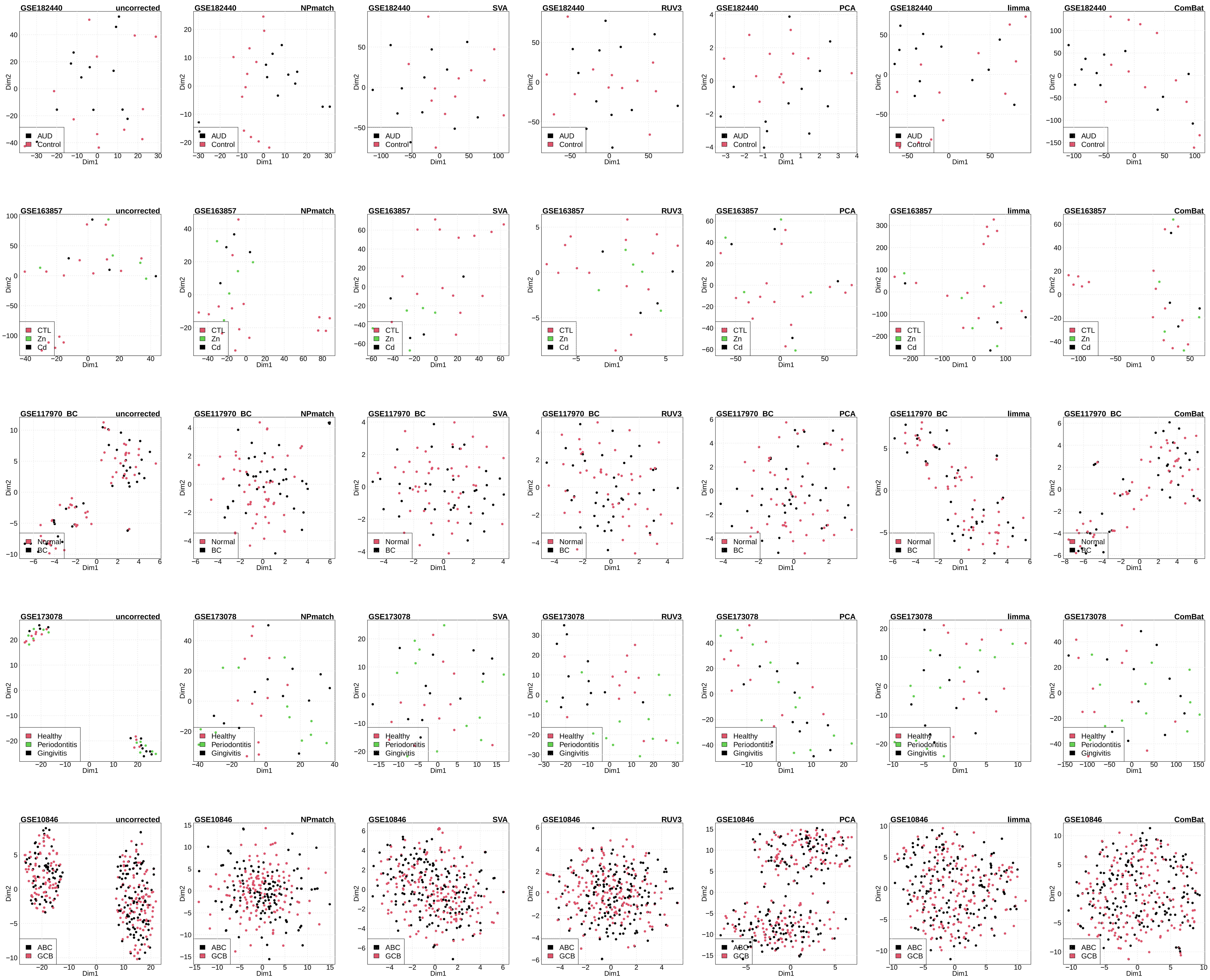

### Figure S4

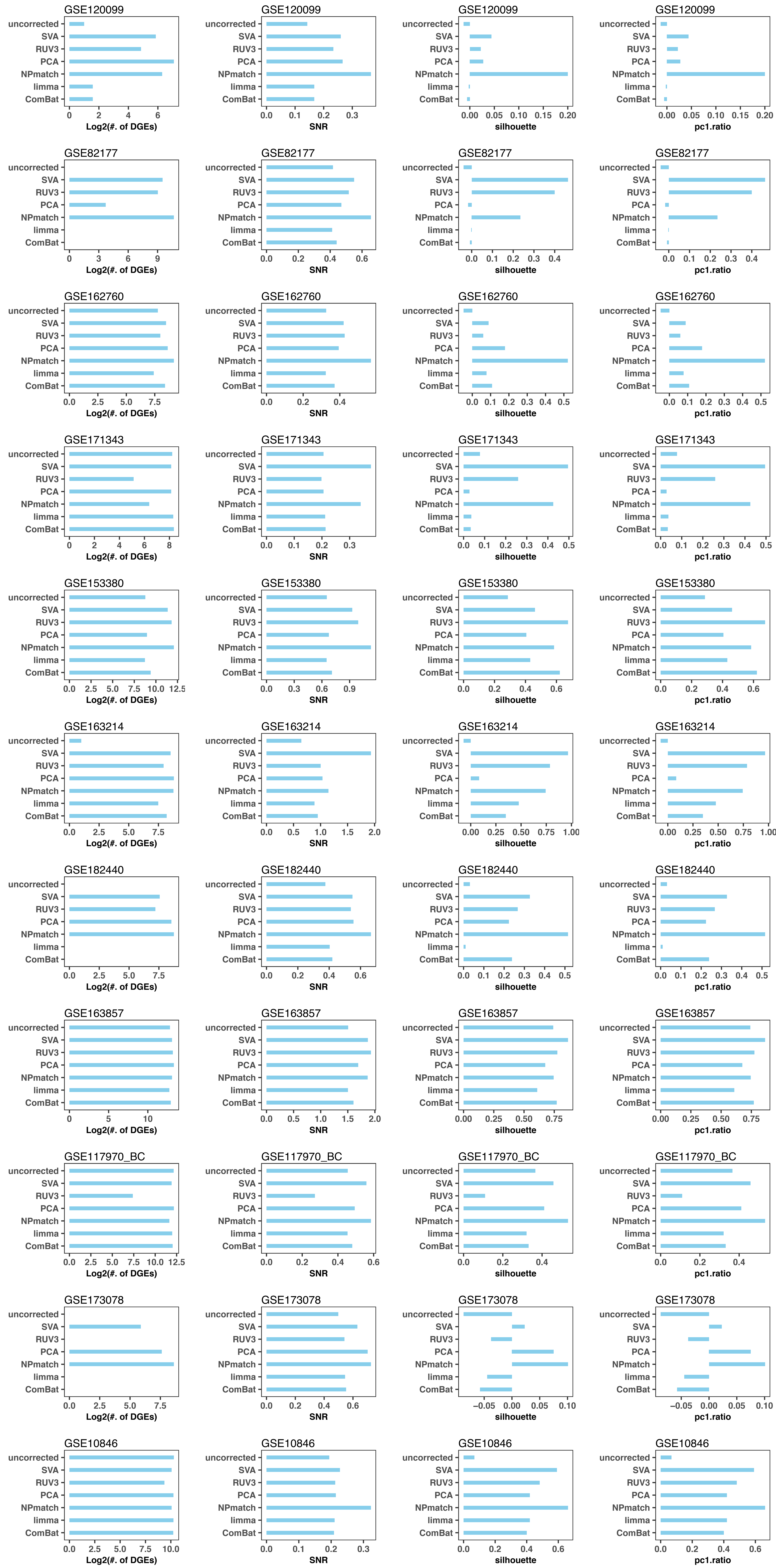
